# A phylogenetic framework of the legume genus *Aeschynomene* for comparative genetic analysis of the Nod-dependent and Nod-independent symbioses

**DOI:** 10.1101/422956

**Authors:** Laurent Brottier, Clémence Chaintreuil, Paul Simion, Céline Scornavacca, Ronan Rivallan, Pierre Mournet, Lionel Moulin, Gwilym P. Lewis, Joël Fardoux, Spencer C. Brown, Mario Gomez-Pacheco, Mickaël Bourges, Catherine Hervouet, Mathieu Gueye, Robin Duponnois, Heriniaina Ramanankierana, Herizo Randriambanona, Hervé Vandrot, Maria Zabaleta, Maitrayee DasGupta, Angélique D’Hont, Eric Giraud, Jean-François Arrighi

## Abstract

**SUMMARY:**

- Some *Aeschynomene* legume species have the property of being nodulated by photosynthetic *Bradyrhizobium* lacking the *nodABC* genes. Knowledge of this unique Nod (factor)-independent symbiosis has been gained from the model *A. evenia* but our understanding remains limited due to the lack of comparative genetics with related taxa using a Nod-dependent process.
- To fill this gap, this study significantly broadened previous taxon sampling, including in allied genera, to construct a comprehensive phylogeny. This backbone tree was matched with data on chromosome number, genome size, low-copy nuclear genes and strengthened by nodulation tests and a comparison of the diploid species.
- The phylogeny delineated five main lineages that all contained diploid species while polyploid groups were clustered in a polytomy and were found to originate from a single paleo-allopolyploid event. In addition, new nodulation behaviours were revealed and Nod-dependent diploid species were shown to be tractable.
- The extended knowledge of the genetics and biology of the different lineages in the legume genus *Aeschynomene* provides a solid research framework. Notably, it enabled the identification of *A. americana* and *A. patula* as the most suitable species to undertake a comparative genetic study of the Nod-independent and Nod-dependent symbioses.

## INTRODUCTION

In the field of nitrogen-fixing symbiosis, scientists have a long-standing interest in the tropical papilionoid legume genus *Aeschynomene* since the discovery of the ability of the species *A. afraspera* to develop abundant nitrogen-fixing stem nodules (Hagerup, 1928). This nodulation behavior is uncommon in legumes, being shared by very few hydrophytic species of the genera *Discolobium, Neptunia* and *Sesbania*, but it is exceptionally widespread among the semi-aquatic *Aeschynomene* species (Alazard, 1985; Boivin *et al*., 1997; Chaintreuil *et al*., 2013). These stem-nodulating *Aeschynomene* species are able to interact with *Bradyrhizobium* strains that display the unusual property of being photosynthetic (Giraud *et al*., 2000; Miché *et al*., 2010). Most outstanding is the evidence that some of these photosynthetic *Bradyrhizobium* strains lack both the *nodABC* genes required for the synthesis of the key “Nod factors” symbiotic signal molecules and a type III secretion system (T3SS) that is known in other rhizobia to activate or modulate nodulation (Giraud *et al*., 2007; Okazaki *et al*., 2013, 2015). These traits revealed the existence of an alternative symbiotic process between rhizobia and legumes that is independent of the Nod factors.

As in the legume genus *Arachis* (peanut), *Aeschynomene* uses an intercellular symbiotic infection process instead of infection thread formation that can be found in other legume groups (Sprent *et al*., 2017). This led to the suggestion that the Nod-independent process might correspond to the ground state of the rhizobial symbiosis, although we cannot exclude that it represents an alternative symbiotic process compared to the one described in other legumes (Sprent & James, 2008; Madsen *et al*., 2010, Okubo *et al*., 2012). It is noteworthy that all the Nod-independent species form a monophyletic clade within the *Aeschynomene* phylogeny and jointly they also display striking differences in the bacteroid differentiation process compared to other *Aeschynomene* species (Chaintreuil *et al*., 2013; Czernic *et al*., 2015). To decipher the molecular mechanisms of this distinct symbiosis, the Nod-independent *A. evenia* has been used as a new model legume, because its genetic and developmental characteristics (diploid with a reasonable genome size −2n=20, 415 Mb/1C-, short perennial and autogamous, can be hybridized and transformed) make this species tractable for molecular genetics (Arrighi *et al*., 2012, 2013, 2015). Functional analyses revealed that some symbiotic determinants identified in other legumes (*SYMRK, CCaMK, HK1* and *DNF1*) are recruited, but several key genes involved in bacterial recognition (e.g. *LYK3*), symbiotic infection (e.g. *EPR3* and *RPG*), and nodule functioning (e.g. *DNF2* and *FEN1*) were found not to be expressed in *A. evenia* roots and nodules, based on RNAseq data (Czernic *et al*., 2015; Fabre *et al*., 2015; Chaintreuil *et al*., 2016a; Nouwen *et al*., 2017). This suggested that the Nod-independent symbiosis is distinct from the Nod-dependent one.

Forward genetics are now expected to allow the identification of the specific molecular determinants of the Nod-independent process in *A. evenia* (Arrighi *et al*., 2012; Chaintreuil *et al*., 2016a). In addition, comparing *A. evenia* with closely related Nod-dependent *Aeschynomene* species will promote our understanding how symbiosis evolved in legumes. The genus *Aeschynomene* (restricted now to the section *Aeschynomene* as discussed in Chaintreuil *et al*. (2013)) is traditionally composed of three infrageneric taxa, subgenus *Aeschynomene* (which includes all the hydrophytic species) and subgenera *Bakerophyton* and *Rueppellia* (Rudd, 1955; Gillet *et al*., 1971). The genus has also been shown to be paraphyletic, with a number of related genera being nested within it, but together they form a distinct clade in the tribe Dalbergieae (Chaintreuil *et al*., 2013; Rudd, 1981; Lavin *et al*., 2001; Klitgaard *et al*., 2005; LPWG, 2017). Within this broad clade, two groups of semi-aquatic *Aeschynomene* have been well-studied from a genetic and genomic standpoint: the *A. evenia* group, which contains all the Nod-independent species (most of them being 2x), and the *A. afraspera* group (all species being Nod-dependent) that appears to have a 4x origin (Arrighi *et al*., 2014, Chaintreuil *et al*. 2016b, 2018). For comparative analyses, the use of Nod-dependent species with a diploid structure would be more appropriate, but such *Aeschynomene* species are poorly documented.

To overcome these limitations, our aim was to produce a species-comprehensive phylogenetic tree supplemented with genetic and nodulation data. For this, we made use of an extensive taxon sampling in both the genus *Aeschynomene* and in closely related genera to capture the full species diversity of the genus and to clarify phylogenetic relationships between taxa. For most species, we also documented chromosome number, genome size and molecular data for low-copy nuclear genes, thus allowing the identification of diploid species as well as untangling the genome structure of polyploid taxa. In addition, these species were characterized for their ability to nodulate with various *Bradyrhizobium* strains containing or lacking *nod* genes and, finally, the diploid species were submitted to a comparative analysis of their properties. In light of the data obtained in this study, we discuss the interest of two *Aeschynomene* species, *A. americana* and *A. patula*, to set up a comparative genetic system to complement the *A. evenia* model.

## MATERIALS & METHODS

### Plant material

All the accessions of *Aeschynomene* used in this study, including their geographic origin and collection data are listed in Tables S1 and S4. Seed germination and plant cultivation in the greenhouse were performed as indicated in Arrighi *et al*. (2012). Phenotypic traits such as the presence of adventitious root primordia and nodules on the stem were directly observed in the glasshouse.

### Nodulation tests

Nodulation tests were carried out using *Bradyrhizobium* strains ORS278 (originally isolated from *A. sensitiva* nodules), ORS285 (originally isolated from *A. afraspera* nodules), ORS285Δ*nod* and DOA9 (originally isolated from *A. americana* nodules) (Giraud *et al*., 2007; Teamtisong *et al*., 2014; Bonaldi *et al*., 2011). *Bradyrhizobium* strains were cultivated at 34°C for seven days in Yeast Mannitol (YM) liquid medium supplemented with an antibiotic when necessary (Howieson *et al*., 2016). Plant *in vitro* culture was performed in tubes filled with buffered nodulation medium (BNM) as described in Arrighi *et al*. (2012). Five-day-old plants were inoculated with 1 mL of bacterial culture with an adjusted OD at 600nm to 1. Twenty one days after inoculation, six plants were analysed for the presence of root nodules. Nitrogen-fixing activity was estimated on the entire plant by measurement of acetylene reducing activity (ARA) and microscopic observations were performed using a stereo-macroscope (Nikon AZ100, Champigny-sur-Marne, France) as published in Bonaldi *et al*. (2011).

### Molecular methods

Plant genomic DNA was isolated from fresh material using the classical CTAB (Cetyl Trimethyl Ammonium Bromide) extraction method. For herbarium material, the method was adapted by increasing the length of the incubation (90 min), centrifugation (20 min) and precipitation (15 min) steps. The nuclear ribosomal internal transcribed spacer region (ITS), the chloroplast *matK* gene and four low-copy nuclear genes (*CYP1, eiF1*α, *SuSy*, and *TIP1;1*) previously identified in the *A. evenia* and *A. afraspera* transcriptomes were used for phylogenetic analyses (Arrighi *et al*., 2014; Chaintreuil *et al*., 2016). The genes were PCR-amplified, cloned and sequenced as described in Arrighi *et al*. (2014) (Table S2). For genomic DNA extracted from herbarium specimens, a battery of primers was developed to amplify the different genes in overlapping fragments as short as 250 bp (Table S2). The DNA sequences generated in this study were deposited in GenBank (Table S3).

### Phylogenetic analyses and traits mapping

Sequences were aligned using MAFFT (*--localpair –maxiterate 1000*; Katoh & Standley, 2013). Phylogenetic reconstructions were performed for each gene as well as for concatenated datasets under a Bayesian approach using Phylobayes 4.1b (Lartillot & Philippe, 2004) and the site-heterogeneous CAT+F81+Γ4 evolution model. For each analysis, two independent chains were run for 10,000 Phylobayes cycles with a 50% burn-in. Ancestral states reconstruction was done through stochastic character mapping using the Phytools R package (Revell, 2012) running 10 simulations for each character.

### Species networks and hybridizations

To test if the phylogeny obtained by concatenating the four low-copy nuclear genes (*CYP1, eiF1*α, *SuSy*, and *TIP1;1*) was most likely obtained by gene duplications followed by differential losses or by a combination of duplications, losses coupled with one or several allopolyploidy events involving *A. patula* and *Soemmeringia semperflorens*, the method presented in To & Scornavacca (2015) was used. In short, this method computes a reconciliation score by comparing a phylogenetic network and one or several gene trees. The method allows allopolyploidy events at hybridization nodes while all other nodes of the network are associated to speciation events; meanwhile, duplication and loss events are allowed at a cost (here, arbitrarily fixed to 1) on all nodes of the gene tree.

Thus, the set of 4 nuclear gene trees was used to score different phylogenetic networks corresponding to four different potential evolutionary histories. Two alternative networks with no reticulation corresponding to the two topologies obtained either with the group A (T1) or group B (T2) served to evaluate a no-allopolyploidisation hypothesis. The topology yielding the best score (T2) served to generate and compare all phylogenetic networks with one or two hybridization nodes, involving *A. patula* and/or *S. semperflorens*, to test successively a one-allopolyploidisation scenario (N1-best) and a two-allopolyploidisation evolutionary scenario (N2-best).

### GBS analysis

A GBS library was constructed based on a described protocol (Oueslati *et al*., 2017). For each sample, a total of 150 ng of genomic DNA was digested using the two-enzyme system, PstI (rare cutter) and Mse (common cutter) (New England Biolabs, Hitchin, UK), by incubating at 37°C for 2 h. The ligation reaction was performed using the T4 DNA ligase enzyme (New England Biolabs, Hitchin, UK) at 22°C for 30 min and the ligase was inactivated at 65°C for 30 min. Ligated samples were pooled and PCR-amplified using the Illumina Primer 1 (barcoded adapter with PstI overhang) and Illumina Primer 2 (common Y-adapter). The library was sequenced on an Illumina HiSeq 3000 (1x 150 pb) (at the Get-PlaGe platform in Toulouse, France).

The raw sequence data were processed in the same way as in the study described in Garsmeur *et al*. (2018). SNP calling from the raw Illumina reads was performed using the custom python pipeline VcfHunter (available at https://github.com/SouthGreenPlatform/VcfHunter/) (Guillaume Martin, CIRAD, France). For all samples, these sequence tags were aligned to the *A. evenia* 1.0 reference genome (JF Arrighi, unpublished data). The SNP results from all the samples were converted into one large file in VCF format and the polymorphism data were subsequently analysed using the web-based application SNiPlay3 (Dereeper *et al*., 2015). First, the SNP data were treated separately for each species and filtered out to remove SNP with more than 10% missing data as well as those with a minor allele frequency (MAF) of 0.01 using integrated VCFtools. Second, an overall representation of the species diversity structures was obtained by making use of the PLINK software as implemented in SNiPlay3. This software is based on the multidimensional-scaling (MSD) method to produce two-dimensional plots. The Illumina HiSeq 3000 sequencing raw data are available in the NCBI SRA (Sequence Read Archive) under the study accession number: SRP149516.

### Genome size estimation and chromosome counting

Genome sizes were measured by flow cytometry using leaf material as described in Arrighi *et al.* (2012). Genome size estimations resulted from measurements of three plants per accession and *Lycopersicum esculentum* (Solanaceae) cv “Roma” (2C = 1.99 pg) was used as the internal standard. The 1C value was calculated and the conversion factor 1 pg DNA = 978 Mb was used to express it in Mb/1C. To count chromosome number, metaphasic chromosomes were prepared from root-tips, spread on slides, stained with 4’,6-diamidino-2-phenylindole (DAPI) and their image captured with a fluorescent microscope as detailed in Arrighi *et al*. (2012).

## RESULTS

### A comprehensive phylogeny of the genus *Aeschynomene* and allied genera

To obtain an in-depth view of the phylogenetic relationships within the genus *Aeschynomene* subgenus *Aeschynomene*, which contains the hydrophytic species, we significantly increased previous sampling levels by the addition of new germplasm accessions and, if these were not available, we used herbarium specimens. DNA was isolated for 40 out of the 41 species (compared to the 27 species used in Chaintreuil *et al.* (2013)) included in this group in taxonomic and genetic studies (Table S1) (Arrighi *et al*., 2014; Chaintreuil *et al*., 2012, 2016b, 2018; Rudd, 1955). In addition, to determine the phylogenetic relationship of this subgenus with respect to *Aeschynomene* subgenera *Bakerophyton* and *Rueppellia*, unclassified *Aeschynomene* species, as well as with the allied genera *Bryaspis, Cyclocarpa, Geissaspis, Humularia, Kotschya, Smithia* and *Soemmeringia*, representatives of these 10 taxa were also sampled (compared to the 5 taxa present in Chaintreuil *et al.* (2013)) (Rudd, 1981; Lewis, 2005). This added 21 species to our total samples (Table S1). The dalbergioid species *Pictetia angustifolia* was used as outgroup (Chaintreuil *et al.* 2013; LPWG, 2017).

Phylogenetic reconstruction of all the taxa sampled was undertaken using Bayesian analysis of the chloroplast *matK* gene and the nuclear ribosomal *ITS* region (Table S2 and S3). The *matK* and *ITS* gene trees distinguished almost all the different *Aeschynomene* groups and related genera (Fig. S1 and S2). The two phylogenetic trees have a very similar topology although some branches of both trees can be lowly supported. Incongruences were also observed for *A. deamii* and the genus *Bryaspis*, but the conflicting placements are poorly supported and were thus interpreted as a lack of resolution typical of single-marker trees, rather than strong incongruence. To improve the phylogenetic resolution among the major lineages, the *matK* gene and the *ITS* sequence datasets were combined into a single phylogenetic analysis where only well-supported nodes were considered (posterior probability (PP) ≥ 0.5) (Fig. 1). Our analysis recovered a grade of five main lineages with a branching order that received robust support (PP≥ 0.92): (1) a basally branching lineage represented by *A. americana*, (2) an *A. montevidensis* lineage, (3) an *A. evenia* lineage corresponding to the Nod-independent clade (Arrighi *et al*., 2012, 2014), (4) a newly-identified lineage containing *A. patula* and (5) a lineage represented by an unresolved polytomy clustering the *A. afraspera* clade (Chaintreuil *et al*., 2016b) with all the remaining taxa.

**Figure 1:**
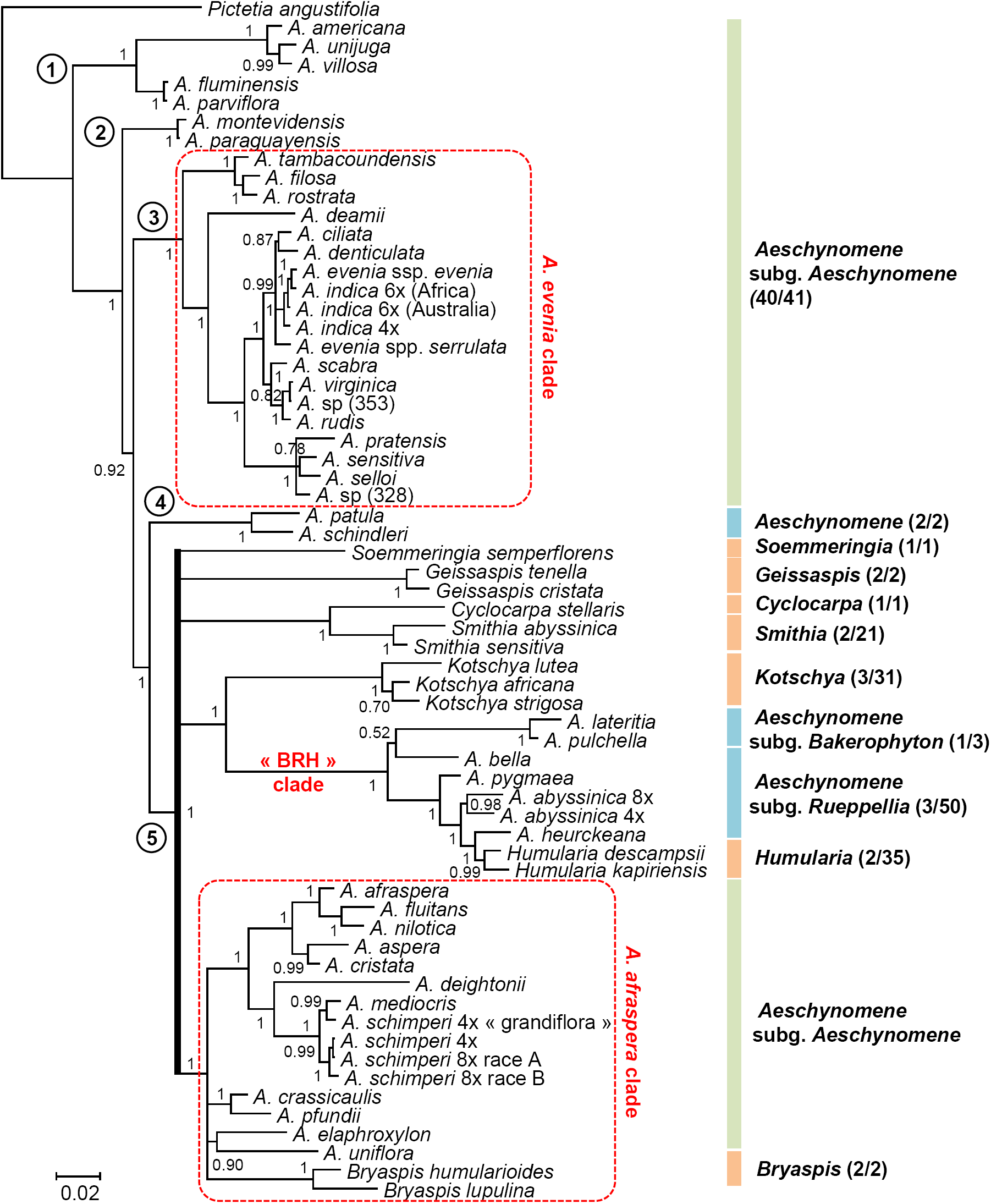
Phylogeny of the genus *Aeschynomene* and allied genera. The Bayesian phylogenetic reconstruction was obtained using the concatenated *ITS* (Internal Transcribed Spacer) **+** *matK* sequences. Numbers at branches indicate posterior probability above 0.5. The five main lineages are identified with a circled number and the two previously studied *Aeschynomene* groups are framed in a red box bordered with a dashed line. On the right are listed *Aeschynomene* subgenus *Aeschynomene* (in green), other *Aeschynomene* subgenera or species groups (in blue) and related genera (in orange).

In large part, our work also provided good species-level resolution and demonstrated that *Aeschynomene* subgenus *Aeschynomene* (as currently circumscribed) is interspersed on the phylogenetic tree with the lineage containing *A. patula*, the two other subgenera of *Aeschynomene* and a number of other genera related to *Aeschynomene* (Fig. 1) (Chaintreuil *et al*., 2013; Lavin *et al*., 2001; LPWG, 2017; Du Puy *et al*., 2002). The combined analysis also grouped the genus *Bryaspis* with the species related to *A. afraspera* in a highly supported clade but its exact position with respect to other taxa remained inconclusive, as previously observed (Fig. 1) (Chaintreuil *et al*., 2013). Most noticeably, several intergeneric relationships are consistently recovered, notably sister-clade relationships between *Cyclocarpa* and *Smithia* as well as between *Aeschynomene* subgenera *Bakerophyton* and *Rueppellia* together with the genus *Humularia* (referred to as the BRH clade herein after) (Fig. 1). This clade supports previous observations of a morphological continuum between *Aeschynomene* subgenus *Rueppellia* and the genus *Humularia* and brings into question their taxonomic separation (Gillett *et al*., 1971).

## Ploidy level of the species and genomic structure of the different lineages

The revised *Aeschynomene* phylogeny was used as a backbone tree to investigate the evolution of ploidy levels. Previous studies had demonstrated that the *A. evenia* clade is mostly diploid (2n=2x=20) even if some species such as *A. indica* (2n=4x=40, 2n=6x=60) appear to be of recent allopolyploid origin (Arrighi *et al*., 2014; Chaintreuil *et al*., 2018). Conversely, all the species of the *A. afraspera* group were found to be polyploid (2n=4x=28,38,40, 2n=8x=56,76) and to have a common AB genome structure but the origin of the polyploidy event remained undetermined (Chaintreuil et al., 2016b). To assess the ploidy levels in *Aeschynomene* species and related genera, chromosome numbers and nuclear DNA content were determined (appended to labels in Fig. 2a, Table S1, Fig. S3 and S4). We provide evidence that the lineages containing *A. americana, A. montevidensis, A. evenia* and *A. patula*, as well as *Soemmeringia semperflorens*, are diploid with 2n=20, with the smallest 2x genome found in *A. patula* (0.58 pg/2C) and the largest 2x genome in *A. deamii* (1.93 pg/2C). With the exception of *S. semperflorens*, all the groups that are part of the polytomy were characterized by higher chromosome numbers 2n=28,36,38,40 (up to 76). These chromosome numbers equate to approximately twice that of the diploid species (except for 2=28), suggesting that the corresponding groups are most probably polyploid. Species with chromosome numbers departing from 2n=40 are likely to be of disploid origin as already described in the *A. afraspera* clade (Chaintreuil *et al*., 2016b). Here again, important genome size variations ranging from 0.71 pg/2C for the *Geissaspis* species to 4.82 pg/2C for the 4x *A. schimperi* highlight the genomic differentiation of the various taxa (Fig. 2a, Table S1).

**Figure 2:**
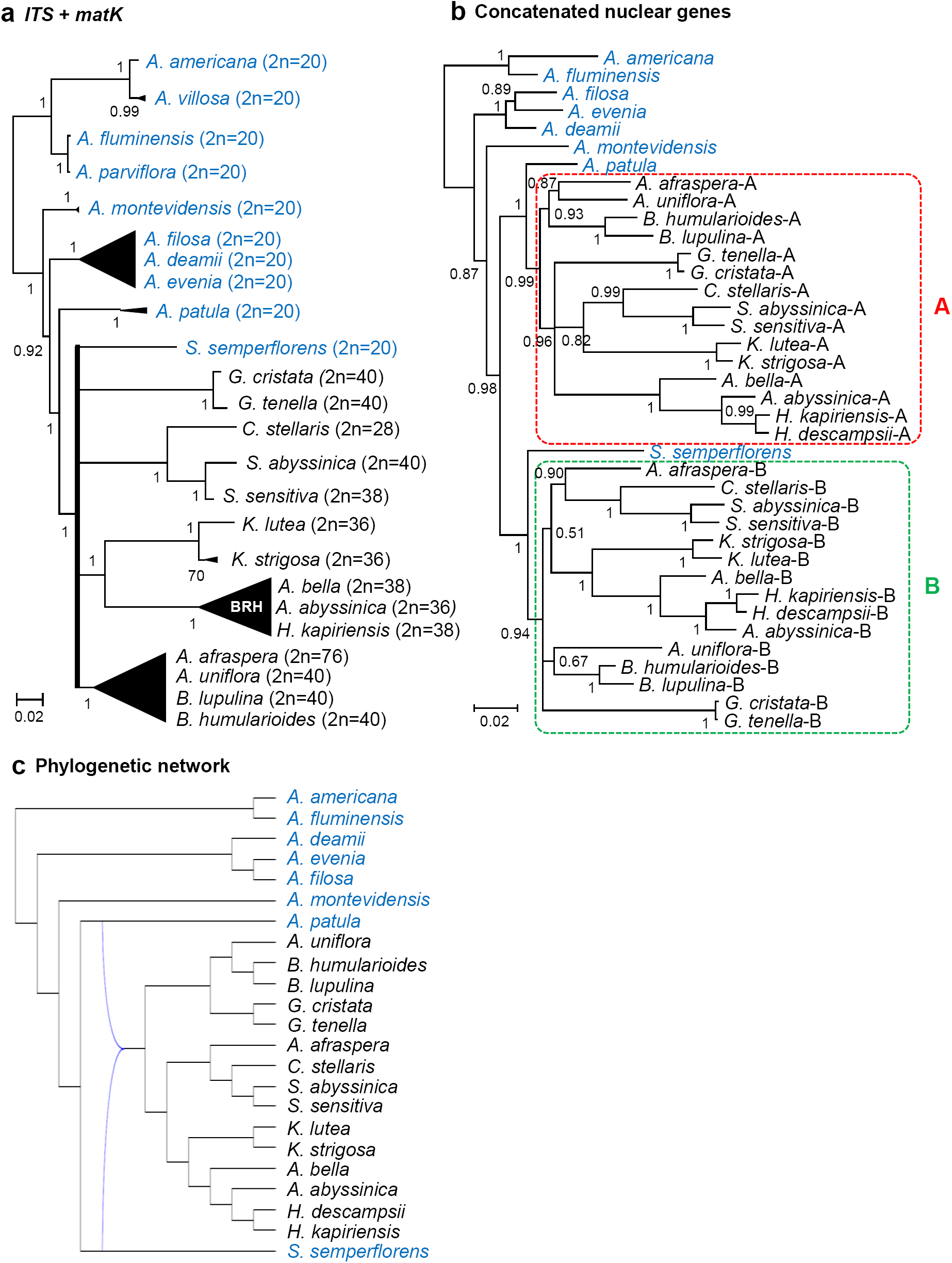
Genomic characteristics and phylogenetic relationships. (a) Simplified Bayesian *ITS*+*matK* phylogeny with representative species of different lineages and groups. The *A. evenia, A. afraspera* and BRH (*Bakerophyton-Rueppelia-Humularia*) clades are represented by black triangles and the polytomy is depicted in bold. Chromosome numbers are indicated in brackets. (b) Phylogenetic relationships based on the combination of 4 concatenated nuclear low-copy genes (*CYP1, eif1a, SuSy* and *TIP1;1* genes detailed in Figure S5). Diploid species (2n=20) are in blue, polyploid species (2n≥28) in black. The A and B subgenomes of the polyploid taxa are delineated by red and green boxes in dashed lines, respectively. Nodes with a posterior probability inferior to 0.5 were collapsed into polytomies. Posterior probability above 0.5 are indicated at every node. (c) The one-allopolyploidisation hypothesis (N1-best) obtained with the phylogenetic network analysis based on the T2 tree with reticulations in blue (detailed in Fig. S7).

To firmly link chromosome numbers to ploidy levels and to clarify genetic relationships between the different lineages, we cloned and sequenced four nuclear-encoded low-copy genes in selected species: *CYP1* (Cyclophilin 1), *eiF1*α (eukaryotic translation initiation factor α), *SuSy* (Sucrose Synthase) and *TIP1;1* (tonoplast intrinsic protein 1;1) (Table S2). For all diploid species, only one gene sequence was obtained, while for all the polyploid species, in almost all cases, a pair of putative homeologues was isolated, thus confirming their genetic status inferred from the karyotypic data (Table S3). In general, the duplicated copies were highly divergent and nested in two different major clades in the resulting Bayesian phylogenic trees generated for each gene (Fig. S5). One clade contained all the A copies (except for one anomalous sequence for *Bryaspis lupulina* in the *eiF1*α tree) and the other clade gathered all the B copies previously identified in *A. afraspera* (Chaintreuil *et al*., 2016b). These two clades A and B did not always receive high support, however it is notable that the A copies formed a monophyletic group with, or sister to, the *A. patula* sequence and similarly the B copies with, or sister to, the *Soemmerignia semperflorens* sequence, in all gene trees (Fig. S5). In an attempt to improve phylogenetic resolution, the four gene data sets were concatenated. This combination resulted in a highly supported Bayesian tree that places the A copy clade as the sister to the diploid *A. patula* (PP =1), and the B copy clade as sister to the diploid *S. semperflorens* (PP =1) (Fig. 2b). As a result, these phylogenetic analyses combined to karyotypic data show that all the five main lineages contain diploid species. They also reveal that all the polyploid groups share the same AB genome structure, with the diploid *A. patula* and *S. semperflorens* species being the closest modern representatives of the ancestral donors of the A and B genomes.

In addition, an ancestral state reconstruction analysis performed on the *ITS*+*matK* phylogeny indicates that diploidy is the ancestral condition in the whole revised group and that tetraploidy most likely evolved once in the polytomy (Fig. S6). To provide support on a probable single origin of the allopolyploidy event, separate and concatenated nuclear gene trees were further used for a phylogenetic network analysis. In this analysis, the two non-allopolyploidisation hypotheses (T1 and T2) were found to be more costly (scores of 207 and 196) than the two hypotheses allowing for hybridization (N1-best and N2-best with scores of 172 and 169, respectively) (Fig. S7a-d). The one-allopolyploidisation hypothesis (N1-best) strongly indicates that a hybridization involving the lineages that contain *A. patula* and *S. semperflorens* gave rise to all the polyploid groups (Fig. S7c). Although the two-allopolyploidisation hypothesis (N2-best) yielded the absolute best score, the score improvement was very low (169 vs 172) and the resulting network included the hybridization inferred with the one-allopolyploidisation hypothesis making this latter hypothesis most probably the correct one (Fig. S7d).

## Nodulation properties of the different *Aeschynomene* lineages

Species of *Aeschynomene* subgenus *Aeschynomene* are known to be predominantly amphibious and more than 15 of these hydrophytic species (found in the *A. evenia* and *A. afraspera* clades, as well as *A. fluminensis*) have been described as having the ability to develop stem nodules (Boivin *et al*., 1997; Chaintreuil *et al*., 2016b; Lock, 1989; Rudd, 1955). In *A. fluminensis*, these nodules are observed only on submerged stems (as also seen in the legume *Discolobium pulchellum*), while they occur on aerial stems within the *A. evenia* and *A. afraspera* clades (Fig. 3a) (Alazard & Duhoux, 1987; Chaintreuil *et al*., 2013; Loureiro *et al*., 1994, 1995). Phenotypic analysis of representatives of the different lineages under study revealed that they all display adventitious root primordia along the stem (Fig. 3a,b). Adventitious roots are considered to be an adaptation to temporary flooding and they also correspond to nodulation sites in stem-nodulating *Aeschynomene* species (Fig. 3b) (Alazard & Duhoux, 1987). Given that the *A. evenia* and *A. afraspera* clades are now demonstrated not to share the same genomic components provides a genetic argument for independent developments of stem nodulation by photosynthetic bradyrhizobia. Reconstruction of ancestral characters based on the *ITS*+*matK* phylogeny confirmed that the whole group was ancestrally a wet ecology taxon endowed with adventitious root primordia but that the stem nodulation ability evolved several times, as previously inferred (Fig. S8, S9; and S10) Chaintreuil *et al*., 2013, 2016b).

**Figure 3:**
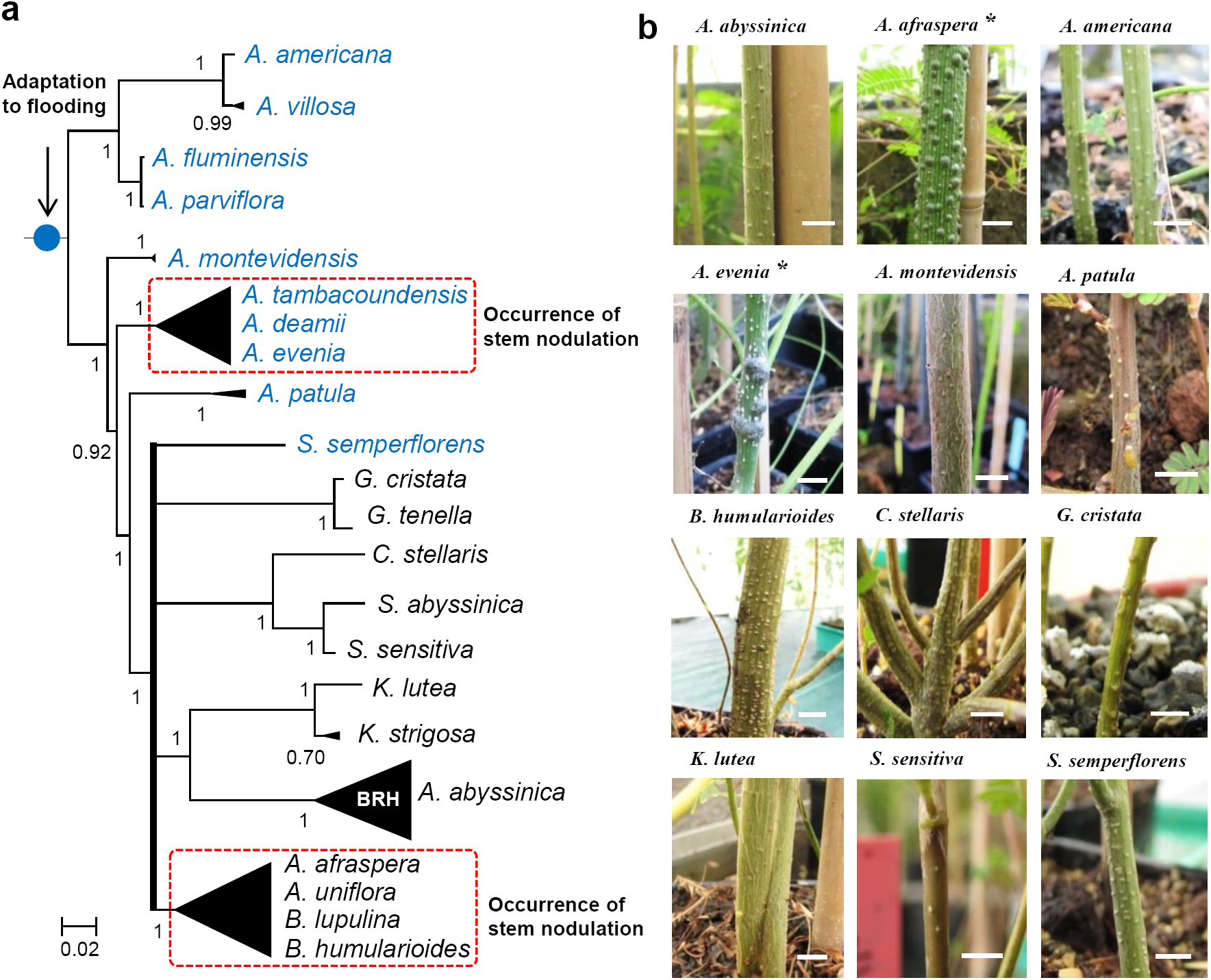
Occurrence of adventitious root primordia and of stem nodulation. (a) Simplified Bayesian *ITS*+*matK* phylogeny of the whole group with the *A. evenia, A. afraspera* and BRH (*Bakerophyton-Rueppelia-Humularia*) clades represented by black triangles. The polytomy is depicted in bold. The shared presence of adventitious root primordia is depicted on the stem by a blue circle. Dashed red boxes indicate groups comprising aerial stem-nodulating species. Asterisks refer to illustrated species in (b) for aerial stem-nodulation. (b) Stems of representatives for the different lineages and groups. Small spots on the stem correspond to dormant adventitious root primordia and stem nodules are visible on the species marked by an asterisk. Bars: 1cm.

To investigate whether the newly studied species could be nodulated by photosynthetic bradyrhizobia, we extended the results obtained by Chaintreuil *et al*. (2013) by testing the nodulation abilities of 22 species available (listed in Fig. 4a) for which adequate seed supply was available. Three different strains of *Bradyrhizobium* equating to the three cross-inoculation (CI) groups defined by Alazard (1985) were used: DOA9 (non-photosynthetic *Bradyrhizobium* of CI-group I), ORS285 (photosynthetic *Bradyrhizobium* with *nod* genes of CI-group II) and ORS278 (photosynthetic *Bradyrhizobium* lacking *nod* genes of CI-group III). These strains were used to inoculate the 22 species and their ability to nodulate them was analysed at 21 dpi. For this, we recorded nodule formation and compared nitrogen fixation efficiency by an acetylene reduction assay (ARA) and observation of plant vigor. Nodulation was observed on all species tested except for *Smithia sensitiva* that had a problem with root development, *A. montevidensis* and *S. semperflorens*. For these three species, either the culture conditions or the *Bradyrhizobium* strains used were not appropriate (Fig. 4a).

**Figure 4:**
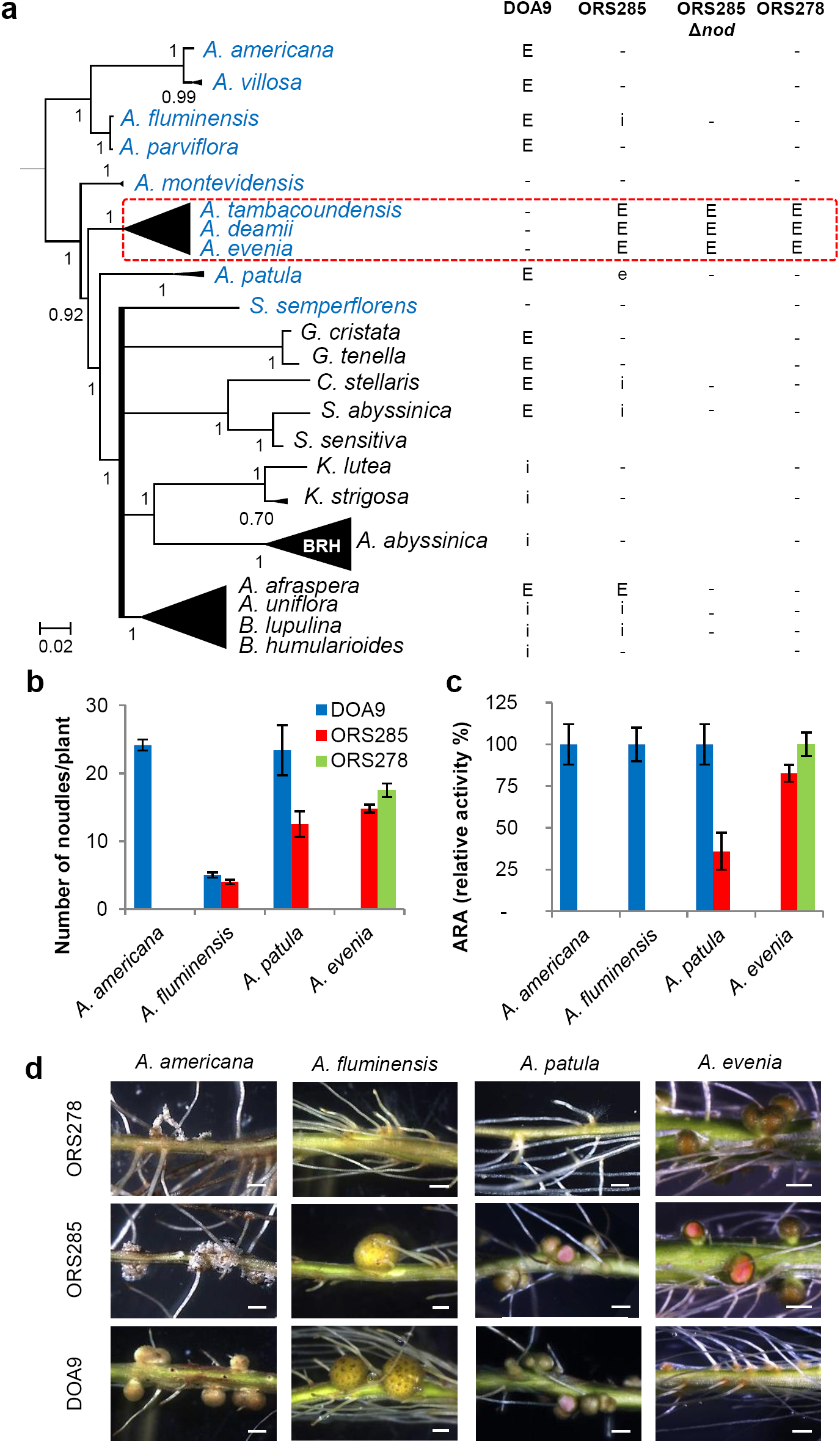
Comparison of the root nodulation properties. (a) Species of different lineages and groups that were tested for nodulation are listed in the simplified Bayesian phylogeny on the left. Root nodulation tests were performed using the DOA9, ORS285, ORS285Δ*nod* and ORS278 strains. E, effective nodulation; e, partially effective nodulation; i, ineffective nodulation, -, no nodulation; blank, not tested. (b) Number of nodules per plant, (c) relative acetylene-reducing activity (ARA) and (d) aspect of the inoculated roots developing nodules or not (some nodules were cut to observe the leghemoglobin color inside) after inoculation with *Bradyrhizobium* DOA9, ORS285 and ORS278 on *A. americana, A. patula, A. afraspera* and *A. evenia*. Error bars in (b) and (c) represent s.d. (n=6). Scale bar in (d): 1 mm.

The non-photosynthetic strain DOA9 displayed a wide host spectrum but was unable to nodulate the Nod-independent species, *A. deamii, A. evenia* and *A. tambacoundensis*. The photosynthetic strain ORS285 efficiently nodulated *A. afraspera* and the Nod-independent *Aeschynomene* species (Fig 4a), as previously reported (Chaintreuil *et al*., 2013). Interestingly, the ORS285 strain was also able to induce nitrogen-fixing nodules in *A. patula* and ineffective nodules were observed on *A. fluminensis* and the genera *Bryaspis, Cyclocarpa* and *Smithia* (Fig. 4a). To examine if in these species the nodulation process relies on a Nod-dependent or Nod-independent symbiotic process, we took advantage of the availability of a Δ*nod* mutant of the strain ORS285. None of them were found to be nodulated by ORS285Δ*nod*, suggesting that the nodule formation depended on a Nod signaling in these species (Fig. 4a). In fact, the ORS285Δ*nod* mutated strain was able to nodulate only species of the *A. evenia* clade, similarly as to the photosynthetic strain ORS278 naturally lacking *nod*- genes (Fig. 4a). Analysis of the evolution of these nodulation abilities by performing an ancestral state reconstruction on the revisited phylogeny indicated several emergences of the ability to interact with photosynthetic bradyrhizobia and a unique emergence of the ability to be nodulated by the *nod* gene-lacking strain as observed earlier (Fig. S11 and Fig. S12) (Chaintreuil *et al*., 2013). From these nodulation tests, different nodulation patterns emerged for the diploid *Aeschynomene* species (as detailed in Fig. 4b-d) with the DOA9 and ORS278 strains being specific to the Nod-dependent and Nod-independent groups respectively and ORS285 showing a gradation of compatibility between both.

## Diversity of the diploid species outside the Nod-independent clade

To further characterize the diploid species that fall outside of the Nod-independent clade, in which *A. evenia* lies, they were analysed for their developmental properties and genetic diversity (Fig. 5a). All species are described as annuals or short-lived perennials (Du Puy *et al*., 2002; Lewis, 2005; Rudd, 1955). As for *A. evenia, A. americana, A. villosa, A. fluminensis, A. parviflora* and *A. montevidensis* are robust and erect, reaching up to 2 m high when mature, whilst *A. patula* and *S. semperflorens* are creeping or decumbent herbs. These differences in plant habit are reflected by the important variation in seed size between these two groups (Fig. 5a). This has an impact on plant manipulation, because for *A. patula* and *S. semperflorens* seed scarification needs to be adapted (25 min with concentrated sulfuric acid instead of 40 min for the other species) and *in vitro* plant growth takes slightly more time to get a root system sufficiently developed for inoculation with *Bradyrhizobium* strains (10 days-post-germination instead of the 5-7 dpi for other species) (Arrighi *et al*., 2012). Consistent flowering and seed production was observed for *A. americana, A. villosa, A. patula* and *S. semperflorens* when grown under full ambient light in the tropical greenhouse in short days conditions as previously described for *A. evenia*, making it possible to develop inbred lines by successive selfing (Fig. 5a) (Arrighi *et al*., 2012). For *A. fluminensis, A. parviflora* and *A. montevidensis*, flowering was sparse or not observed, indicating that favorable conditions for controlled seed set were not met (Fig. 5a).

**Figure 5:**
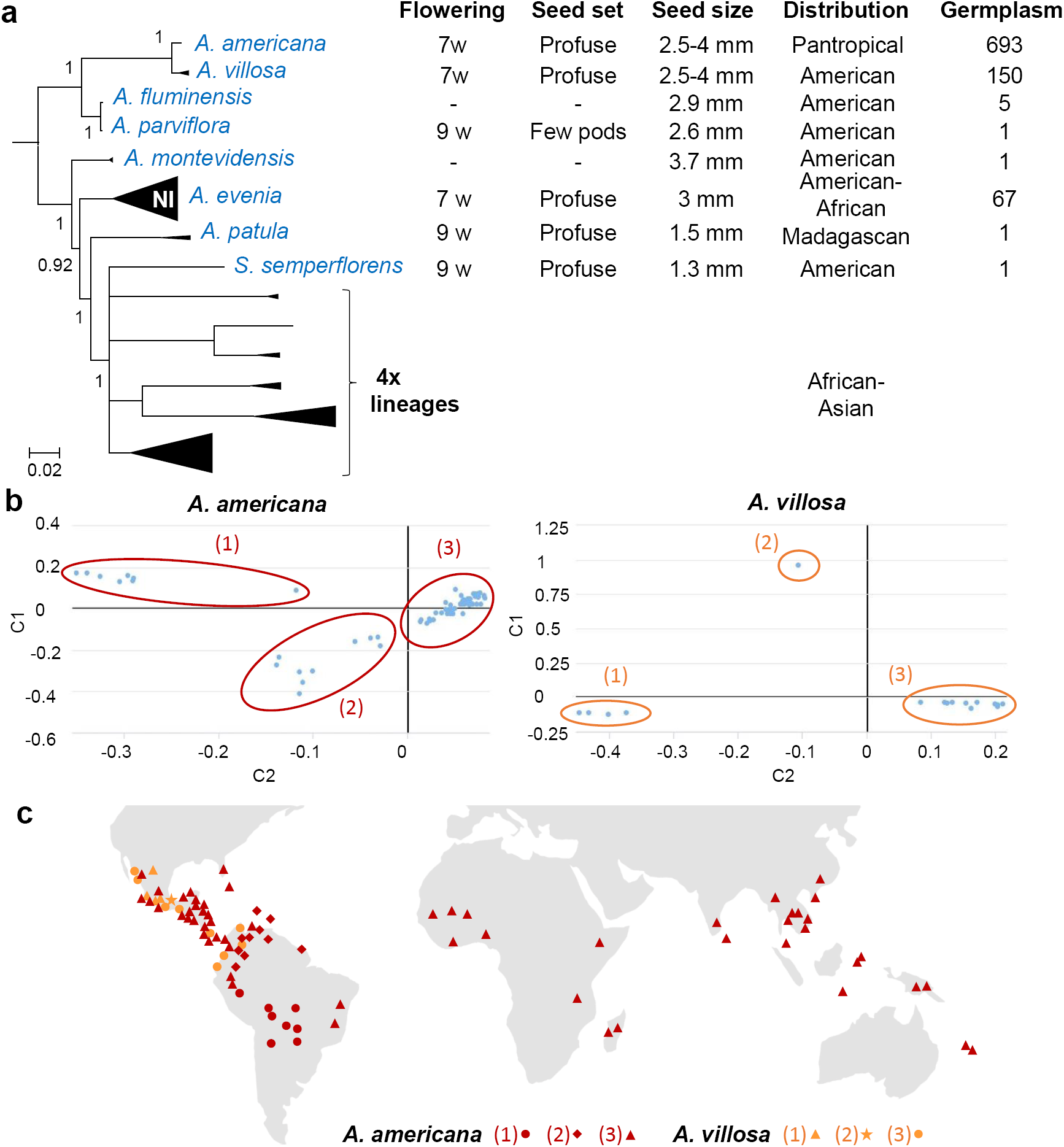
Characteristics of diploid species. (a) Development and germplasm data for species that are listed in the simplified phylogeny on the left. *A. evenia* from the Nod-independent clade (NI) is also included for comparison. Germplasm numbers correspond to the sum of accessions available at CIAT, USDA, Royal Botanic Gardens (Kew), AusPGRIS, IRRI and at LSTM. (b) Multi-dimensional scaling (MSD) plots of the genetic diversity among *A. americana* (left) and *A. villosa* (right) accessions according to coordinates 1 and 2 (C1, C2). Identified groups are delimited by circles and labeled with numbers. (c) Geographical distribution of the of *the A. americana* and *A. villosa* accessions. Taxon colours and group numbers are the same as in (b). Details of the accessions are provided in Table S4.

Five species (*A. villosa, A. fluminensis, A. parviflora, A. montevidensis* and *S*. *semperflorens*) are strictly American while *A. americana* is a pantropical species and *A. patula* is endemic to Madagascar (Du Puy *et al*., 2002; Lock, 1989; Rudd, 1955). Several species have a narrow geographic distribution or seem to be infrequent, explaining the very limited accession availability in seedbanks (Fig. 5a). This is in sharp contrast with both *A. americana* and *A. villosa* that are well-collected, being widely found as weedy plants and sometimes used as component of pasture for cattle (Fig. 5a) (Cook *et al*., 2005). To assess the genetic diversity of these two species, a germplasm collection containing 79 accessions for *A. americana* and 16 accessions for *A. villosa*, and spanning their known distribution was used (Table S4). A Genotyping-By-Sequencing (GBS) approach resulted in 6370 and 1488 high quality polymorphic SNP markers for *A. americana* and *A. villosa* accessions, respectively. These two SNP datasets subsequently served for a clustering analysis based on the multidimensional-scaling (MSD) method. The MSD analysis distinguished three major groups of accessions for both *A. americana* and *A. villosa* along coordinate axes 1 and 2 (Fig. 5b). When mapping the accessions globally, the three groups identified for *A. villosa* were observed together in Mexico and only group (3) extended to the northern part of South America (Fig. 5c, Table S4). Converly, a clear geographical division was observed for *A. americana* with group (1) occupying the central part of South America, group (2) being found in the Caribbean area while group (3) was present in distinct regions from Mexico to Brazil and in across the Palaeotropics (Fig. 5c, Table S4). *A. americana* is hypothesized to be native in America and naturalized elsewhere (Cook *et al*., 2005). The observed distributions in combination with the MSD analysis, accessions being tightly clustered in group (3) compared to groups (1) and (2), support this idea and indicate that group (3) recently spread worldwide.

## DISCUSSION

### A well-documented phylogenetic framework for the legume genus *Aeschynomene*

We produced a new and comprehensive phylogeny of the genus *Aeschynomene* and its closely related genera complemented by gene data sets, genome sizes, karyotypes and nodulation assays. For plant genera, there are few for which documentation of taxonomic diversity is so extensive and supported by a well-resolved, robustly supported phylogeny which reveals the evolutionary history of the group (Govindarajulu *et al*., 2011). Here, the whole group, which includes the genus *Aeschynomene* with its 3 subgenera and its 7 allied genera, is shown to have experienced cladogenesis leading to five main lineages, including the Nod-independent clade, with diploid species found in all these lineages. The multigene data analysis provided robust evidence that two of them, represented by the two diploid species *A. patula* and *S. semperflorens*, are involved in an ancient allotetraploidization process that gave rise to the different polyploid lineages clustering in a polytomy. Separate allopolyploidization events from the same diploid parents or a single allopolyploid origin are plausible explanations for the formation of these lineages. However, the consistent resolution of the phylogenetic tree obtained with the combined gene data, where *A. patula* and *S. semperflorens* are sisters to the A and B subgenomic sequences, favours the hypothesis of a single allopolyploid origin, as also argued for other ancient plant allopolyploid events in *Asimitellaria* (Saxifragaceae) and *Leucaena* (Leguminosae) (Govindarajulu *et al*., 2011; Okuyama *et al*., 2012). The phylogenetic network analysis also supports the one-allopolyploidisation hypothesis. However, additional nuclear genes will be needed to conclusively confirm that no additional hybridization event occurred. Although not the focus of the present study, it is worth noting that most diploid species are found in the Neotropics, the two modern representatives of the A and B genome donors that gave rise to the 4x lineages are located on different continents (*S. semperflorens* in South America and *A. patula* in Madagascar) and that all the 4x lineages are located in the Palaeotropics (Lewis *et al*., 2005). This raises questions about the evolution of the whole group and the origin of the 4x lineages. In addition, the presence of a polytomy suggests that this allopolyploid event preceded a rapid and major diversification of 4x groups that have been ascribed to different *Aeschynomene* subgenera or totally distinct genera that altogether represent more than 80% of the total species of the whole group (LPWG, 2017; Whitefiel & Lockhart, 2007). Diversification by allopolyploidy occurred repeatedly in the genus *Aeschynomene* since several neopolyploid species are found in both the *A. evenia* clade and the *A. afraspera* clade as exemplified by *A. indica* (4x, 6x) and *A. afraspera* (8x) (Arrighi *et al*., 2014; Chaintreuil *et al*., 2016b). Dense sampling for several *Aeschynomene* taxa or clades also a more precise delimitation of species boundaries (for morphologically similar taxa but which are genetically differentiated or correspond to different cytotypes) and revealed intraspecific genetic diversity that is often geographically-based as showed for the pantropical species *A. americana* (this study), *A. evenia, A. indica* and *A. sensitiva* (Chaintreuil *et al*., 2018). All these *Aeschynomene* species share the presence of adventitious root primordia on the stem that correspond to the infection sites for nodulation. The consistent presence of adventitious root primordia in all taxa of the whole group, together with an ancestral state reconstruction, substantiates the two-step model proposed earlier for the evolution of stem nodulation in *Aeschynomene*, with a common genetic predisposition at the base of the whole group to produce adventitious root primordia on the stem, as an adaptation to flooding, and subsequent mutations occurring independently in various clades to enable stem nodulation (Chaintreuil *et al*., 2013). The ability to interact with photosynthetic bradyrhizobia that are present in aquatic environments also appears to have evolved several times. This photosynthetic activity is important for the bacterial symbiotic lifestyle as it provides energy usable for infection and subsequently for nitrogenase activity inside the stem nodules (Giraud *et al*., 2000). To date, natural occurrence of nodulation by photosynthetic bradyrhizobia has been reported only for the *A. evenia* and *A. afraspera* clades, and for *A. fluminensis* (Loureiro *et al*., 1995; Miché *et al*., 2010; Molouba *et al*., 1997). Nevertheless, we could not test the photosynthetic strains isolated from *A. fluminensis* nodules and the nature of the strains present in those of the newly studied species *A. patula* has not been investigated yet. They would allow the comparison of their nodulation efficiency with the reference photosynthetic *Bradyrhizobium* ORS278 and ORS285 strains. In addition, we can ask if the semi-aquatic lifestyle and/or nodulation with photosynthetic bradyrhizobia may have facilitated the emergence of the Nod-independent symbiosis in the *A. evenia* clade.

### *Aeschynomene* species for a comparative analysis of nodulation with *A. evenia*

To uncover whether the absence of detection for several key symbiotic genes in the root and nodule transcriptomic data of *A. evenia* are due to gene loss or extinction, and to identify the specific symbiotic determinants of the Nod-independent symbiosis, a genome sequencing combined with a mutagenesis approach is presently being undertaken for *A. evenia* in our laboratory. A comparative analysis with Nod-dependent *Aeschynomene* species is expected to consolidate this genomic and genetic analysis performed in *A. evenia* by contributing to elucidate the genetic changes that enabled the emergence of the Nod-independent process. Phylogenomics and comparative transcriptomics, coupled with functional analysis, are undergoing increased development in the study of symbiosis. They enabled unravelling gene loss linked to the lack of developing a symbiosis in certain plant lineages but also to identify new symbiosis genes (for arbuscular mycorrhizal symbiosis (Delaux *et al*., 2014; Bravo *et al*., 2016); for the nodulating symbiosis (Delaux *et al*., 2015; Griesmann *et al*., 2018). Comparative work on symbiotic plants is often hindered, however, either by the absence of closely related species which display gain or loss of symbiotic function or, when these are present, by the lack of a well-understood genetic framework, as outlined in Behm *et al*. (2014), Delaux *et al*. (2015), Geurts *et al*. (2016), Sprent (2017). Nevertheless, in the case of the nodulating *Parasponia*/non-nodulating *Trema* system, a fine comparative analysis was very powerful to demonstrate a parallel loss of the key symbiotic genes *NFR5, NIN* and *RGP,* in the non-nodulating species, challenging the long-standing assumption that *Parasponia* specifically acquired the potential to nodulate (Behm *et al*., 2014; Geurts *et al*., 2016; van Velzen *et al*., 2018). In this respect, the uncovering of the genetic evolution of the genus *Aeschynomene* and related genera along with the identification of diploid species outside of the Nod-independent clade, provided a robust phylogenetic framework that can now be exploited to guide the choice of Nod-dependent diploid species for comparative genetic research. Among these, some species are discarded because of the lack of nodulation with reference *Bradyrhizobium* strains or the inability to produce seeds under greenhouse conditions. Based on efficient nodulation, short flowering time and ease of seed production, *A. americana* (2n=20, 600 Mb) and *A. patula* (2n=20, 270 Mb) appear to be the most promising Nod-dependent diploid species to develop a comparative genetic system with *A. evenia* (2n=20, 400 Mb). In contrast to *A. evenia, A. americana* is nodulated only by non-photosynthetic bradyrhizobia, and in this respect, it behaves in a similar way to other legumes. This species is widespread in the tropics, so that adequate quantities of germplasm are available, and it has already been subject to detailed studies, notably to isolate its nodulating *Bradyrhizobium* strains, among which is the DOA9 strain (Noisangiam *et al*., 2012; Teamtisong *et al*., 2014). As *A. americana* belongs to the most basal lineage in the *Aeschynomene* phylogeny, it may be representative of the ancestral symbiotic mechanisms found in the genus. On the other hand, *A. patula* has a restricted Madagascan distribution with only one accession being available, but it is of interest due to its relatively smaller plant size and genome size (actually the smallest diploid genome in the group) making this species the “Arabidopsis” of the genus *Aeschynomene*. As for *A. americana*, this species is efficiently nodulated by non-photosynthetic bradyrhizobia, but it is also compatible with the photosynthetic *nod* gene-containing ORS285 strain. This property makes *A. patula* particularly interesting as it allows direct comparisons of mechanisms and pathways between it and *A. evenia* without the problem of potential strain effects on symbiotic responses. In addition, when considering the *Aeschynomene* phylogeny, *A. patula* is more closely related to *A. evenia* than is *A. americana*, and so it may be more suitable to demonstrate the changes necessary to switch a Nod-dependent to a Nod-independent process or vice-versa. Developing sequence resources and functional tools for *A. americana* and/or *A. patula* is now necessary to set up a fully workable comparative *Aeschynomene* system. In the long run, handling such a genetic system will be instrumental in understanding how photosynthetic *Bradyrhizobium* and some *Aeschynomene* species co-evolved and in unravelling the molecular mechanisms of the Nod-independent symbiosis.

## ACKNOWLEDGEMENTS

We thank the different seed banks and herbaria for provision of seeds and herbarium vouchers that were used in this study. The present work has benefited from the facilities and expertise of the cytometry facilities of Imagerie-Gif (http://www.i2bc.paris-saclay.fr/spip.php?article279) and of the molecular cytogenetic facilities of the AGAP laboratory (http://umr-agap.cirad.fr/en/plateformes/plateau-de-cytogenetique-moleculaire). This work was supported by a grant from the French National Research Agency (ANR-AeschyNod-14-CE19-0005-01) that financed the design of the study, experimentation and analysis of the data.

## AUTHOR CONTRIBUTIONS

J.F.A. designed the experiments. L.B., C.C., R.R., J.F., M.G.M, S.C.B., C.H., M.D. and J.F.A. performed the experiments and obtained the data. P.S., C.S., L.M. undertook the phylogenetic analyses. P.M., J.Q., G.P.L., X.P., A.D’H., E.G. and J.F.A. analysed the data. M.G., R.D., H. Randriambanona, H. Ramanankierna, H.V. and M.Z. contributed to the acquisition and analysis of accessions. J.F.A. wrote the paper. L.B. and C.C contributed equally. All authors read and approved the final manuscript.

---

**Figure S1: *matK* phylogeny of the genus *Aeschynomene* and allied genera.** Bayesian phylogenetic reconstruction obtained using the chloroplast *matK* gene. Numbers at branches are posterior probability.

**Figure S2: *ITS* phylogeny of the genus *Aeschynomene* and allied genera.** Bayesian phylogenetic reconstruction obtained using the Internal Transcribed Spacer (*ITS*) sequence. Numbers at branches are posterior probability.

**Figure S3: Chromosome numbers in *Aeschynomene* species.** Root tip metaphase chromosomes stained in blue with DAPI (4’,6-diamidino-2-phenylindole). Chromosome numbers are indicated in brackets. Scale bars: 5 µm.

**Figure S4: Chromosome numbers in species of genera related to*Aeschynomene*** Root tip metaphase chromosomes stained in blue with DAPI (4’,6-diamidino-2-phenylindole). Chromosome counts are indicated in brackets. Scale bars: 5 µm.

**Figure S5: Phylogenetic trees based on nuclear low-copy genes.** Bayesian phylogenetic reconstructions obtained for the *CYP1, eif1a, SuSy* and *TIP1;1* genes. Diploid species (2n=20) are in blue, polyploid species (2n≥28) in black excepted *A. afraspera* for which the A and B gene copies are distinguished in red and green respectively. -A, -A1, - A2, -B, -B1 and -B2 indicated the different copies found. Putative A and B subgenomes of the polyploid taxa are delineated by red and green boxes in dashed lines, respectively. Numbers at branches represent posterior probability.

**Figure S6: Ancestral state reconstruction of ploidy levels in the genus *Aeschynomene* and allied genera.** Ancestral state reconstruction was estimated in SIMMAP software using the 50% majority-rule topology obtained by Bayesian analysis of the combined *ITS*+*matK* sequences. Ploidy levels are indicated by different colors. Unknown ploidy levels are denoted by a dash.

**Figure S7: Phylogenetic networks based on the four nuclear *CYP1, eif1a, SuSy* and *TIP1;1* genes.** (a) No-allopolyploidisation hypothesis (T1) based on the concatenated gene tree obtained taking into account the group A (Fig. 2b). (b) No-allopolyploidisation hypothesis (T2) based on the concatenated gene tree obtained taking into account the group B (Fig. 2b). (c) One-allopolyploidisation hypothesis (N1-best). (d) Two-allopolyploidisation hypothesis (N2-best). Blue lines indicate reticulations while other nodes of the network are associated to speciation events. Reconciliation scores obtained for each phylogenetic network are indicated.

**Figure S8: Ancestral state reconstruction of adventive root primordia in the genus *Aeschynomene* and allied genera.** Ancestral state reconstruction was estimated in SIMMAP software using the 50% majority-rule topology obtained by Bayesian analysis of the combined *ITS*+*matK* sequences. Data on the adventitious root primordia come from the present analysis and pertinent previously published data. Presence or not of adventitious root primordia is indicated by different colors.

**Figure S9: Ancestral state reconstruction of ecological habit in the genus *Aeschynomene* and allied genera.** Ancestral state reconstruction was estimated in SIMMAP software using the 50% majority-rule topology obtained by Bayesian analysis of the combined *ITS*+*matK* sequences. Data on the species ecology come from pertinent previously published data. Ecological habits are indicated by different colors.

**Figure S10: Ancestral state reconstruction of the aerial stem nodulation ability in the genus *Aeschynomene* and allied genera.** Ancestral state reconstruction was estimated in SIMMAP software using the 50% majority-rule topology obtained by Bayesian analysis of the combined *ITS*+*matK* sequences. Data on the occurrence of stem nodulation come from pertinent previously published data. Occurrence or not of stem nodulation is indicated by different colors.

**Figure S11: Ancestral state reconstruction of the ability to nodulate with the photosynthetic *Bradyrhizobium* strains in the genus *Aeschynomene* and allied genera.** Ancestral state reconstruction was estimated in SIMMAP software using the 50% majority-rule topology obtained by Bayesian analysis of the combined *ITS*+*matK* sequences. Data on nodulation with photosynthetic *Bradyrhizobium* strains come from the present analysis and pertinent previously published data. Nodulation with photosynthetic *Bradyrhizobium* strains is considered positive only if reported as occurring naturally or being efficient *in vitro*.

**Figure S12: Ancestral state reconstruction of the ability to nodulate with the photosynthetic *Bradyrhizobium* strain ORS278 in the genus *Aeschynomene* and allied genera.** Ancestral state reconstruction was estimated in SIMMAP software using the 50% majority-rule topology obtained by Bayesian analysis of the combined *ITS*+*matK* sequences. Data on nodulation with ORS278 come from the present analysis and pertinent previously published data. Ability or not to nodulate with ORS278 is indicated by different colors.

